# *In vivo* crosslinking and effective 2D enrichment for interactome studies of the nucleosome

**DOI:** 10.1101/2025.02.25.640081

**Authors:** Philipp Bräuer, Laszlo Tirian, Fränze Müller, Karl Mechtler, Manuel Matzinger

**Affiliations:** Research Institute of Molecular Pathology (IMP), Vienna BioCenter (VBC), Vienna, Austria; Gregor Mendel Institute of Molecular Plant Biology (GMI), Austrian Academy of Sciences, Vienna BioCenter (VBC), Vienna, Austria; Institute of Molecular Biotechnology (IMBA), Austrian Academy of Sciences, Vienna BioCenter (VBC), Vienna, Austria

**Keywords:** XL-MS, DSBSO, affinity enrichment, DDX39B

## Abstract

Cross-linking mass spectrometry has evolved as a powerful technique to study protein-protein interactions and to provide structural information over the past decades. Low reaction efficiencies, and complex matrices lead to challenging system wide crosslink analysis. In this study, we improved and streamlined an Azide-A-DSBSO based *in vivo* crosslinking workflow employing two orthogonal effective enrichment steps: Affinity enrichment and size exclusion chromatography (SEC). Combined, they allow an effective pulling of DSBSO containing peptides and remove the background of linear as well as mono-linked peptides. We found that the analysis of a single SEC fraction is effective to yield ∼90% of all crosslinks, which is important whenever measurement time is limited, and sample throughput is crucial. Our workflow resulted in more than 5000 crosslinks from K562 cells and generated a comprehensive PPI network from whole cells as well as nuclear extracts. From 393 PPI found within the nucleus, 56 have not yet been reported in the STING database and are representing a valuable resource for investigating new molecular mechanisms and provide complementary data for future studies. We further show, that by applying DSBSO to nuclear extracts we yield more crosslinks on lower abundant proteins and showcase this on the DEAD-box RNA helicase DDX39B which is predominantly expressed in the nucleus. Our data indicates that DDX39B is present in monomeric and dimeric form together with DDX39A within the nuclear extracts analyzed.

## INTRODUCTION

Protein-protein interactions (PPIs) are fundamental to functional networks and biological processes. Studying the structural and dynamic organization of protein interaction networks in their native cellular environment is challenging. Traditional methods like co-immunoprecipitation (Co-IP), yeast two-hybrid (Y2H), and proximity-labeling mass spectrometry (MS) might be limited by antibody availability and require cell lysis, which leads to a loss of the native environment and limits the detection of transient or weak interactions. High-resolution three-dimensional (3D) protein structure information is typically obtained by techniques such as X-ray crystallography, nuclear magnetic resonance spectroscopy, or cryo-electron microscopy, which require large amounts of purified proteins, again removing protein complexes from their native environment.

Chemical cross-linking mass spectrometry (XL-MS) has emerged as a complementary tool, providing low-resolution native structure information.^1^ Recent advancements in cross-linker (XL) molecules, enrichment strategies, MS methods, and data interpretation software have highlighted the growing importance of XL-MS.^2^

XL-MS can identify PPIs on a proteome-wide level in their native environment and is therefore of great importance for studying biological processes such as cellular signaling pathways and transient protein interactions. However, proteome-wide studies pose a challenge for data analysis (n^2^ problem^3^) and have to deal with high sample complexity, dynamic abundance ranges, and low cross-linking efficiencies. To overcome these challenges the development of MS-cleavable XLs, which contain labile bonds that cleave upon collisional activation to produce characteristic signature ions, has significantly advanced the field.^4^ In situ applications face additional difficulties, such as cell membrane penetration and high hydrolysis rates of XL molecules further reducing reaction efficacy.

To address these challenges, we use disuccinimidyl bis-sulfoxide (DSBSO), a MS cleavable, enrichable linker that was shown to be surprisingly well membrane permeable^5^ and that was already successfully used for *in vivo* studies before.^6,7^ We applied a streamlined sample preparation protocol improving previously reported protocols^8,9^ using complementary enrichment strategies to investigate the nuclear interactome in K562 cells. We further investigate interaction partners and align our experimentally found crosslinks to the predicted 3D structure of DDX39B. This DEAD-box RNA helicase, also known as UAP56, is primarily localized in the nucleus and plays a crucial role in pre-mRNA splicing, mRNA export, and genome stability maintenance. It is involved in a multitude of essential cellular processes and its dysregulation is linked to various cancers, making it a potential target for therapeutic interventions.^10,11^

## RESULTS

### Optimizing enrichment workflow

To capitalize on the advantages of an azide tagged linker such as DSBSO we directly enriched cross-linked peptides on alkyne functionalized beads. Since monolinked peptides, where one end of the crosslinker molecule is connected to a peptide while the other end was hydrolyzed in an aqueous environment, are co-enriched, a second – orthogonal enrichment based on size exclusion was applied. We thereby adapted and streamlined **a**. our own previous work^8^, and **b**. a recent protocol from the Lan Huang lab^9^ (Figure 7 /Methods).

For this we first applied DSBSO (2 mM) to the frozen K562 cells, followed by lysis and filter aided sample preparation (FASP). This was followed by the affinity enrichment step (**a**). Thus, we employed a copper free strain-promoted alkyne azide cycloaddition click reaction using dibenzocyclooctin (DBCO) to form a stable triazole covalently connecting crosslinked peptides to the bead material. Instead of using sepharose based bead material, as previously published^8^, we coupled the DBCO group to magnetic beads which allows for facilitated, more effective washing steps yielding improved crosslink numbers. (Supplementary Figure 1). We additionally decided to benchmark magnetic bead material from 2 different vendors: Cytivia and CubeBiotech. Both are functionalized with active N-hydroxy succinimide (NHS) ester groups which we coupled to DBCO-Amine. Their density of active group ranges from 8-14 μmol/mL for the Cytivia to > 15 μmol/ml for the Cube Biotech beads as specified by the manufacturers. To benchmark them for the purpose of affinity enrichment of DSBSO linked peptides we chose purified, DSBSO linked, and digested, Cas9 as model system. We spiked the crosslinked Cas9 peptides into a background of non-crosslinked HeLa peptides in a 1:1 (w:w) ratio, to mimic a more complex sample matrix. Our results showed that the average number of unique crosslink sites (XLs) was improved by >340% showing that coupling to DBCO and enrichment per se worked as expected (Figure 1A). Beads from both vendors performed at a very comparable level with no significant difference by means of identified crosslink sequence matches (CSMs) or co-enriched monolinks (Figure 1B). However, the background of non-crosslinked, linear peptides that bound unspecifically seemed to be slightly lowered for beads from Cube Biotech (Figure 1C). The Cube Biotech beads yielded less reproducible results though (Figure 1A-C) and were less convenient to handle as they were quite sticky. In contrast, the Cytivia beads demonstrated a non-adherent behavior eliminating smearing effects and the need to use any detergents.

**Figure 1:**
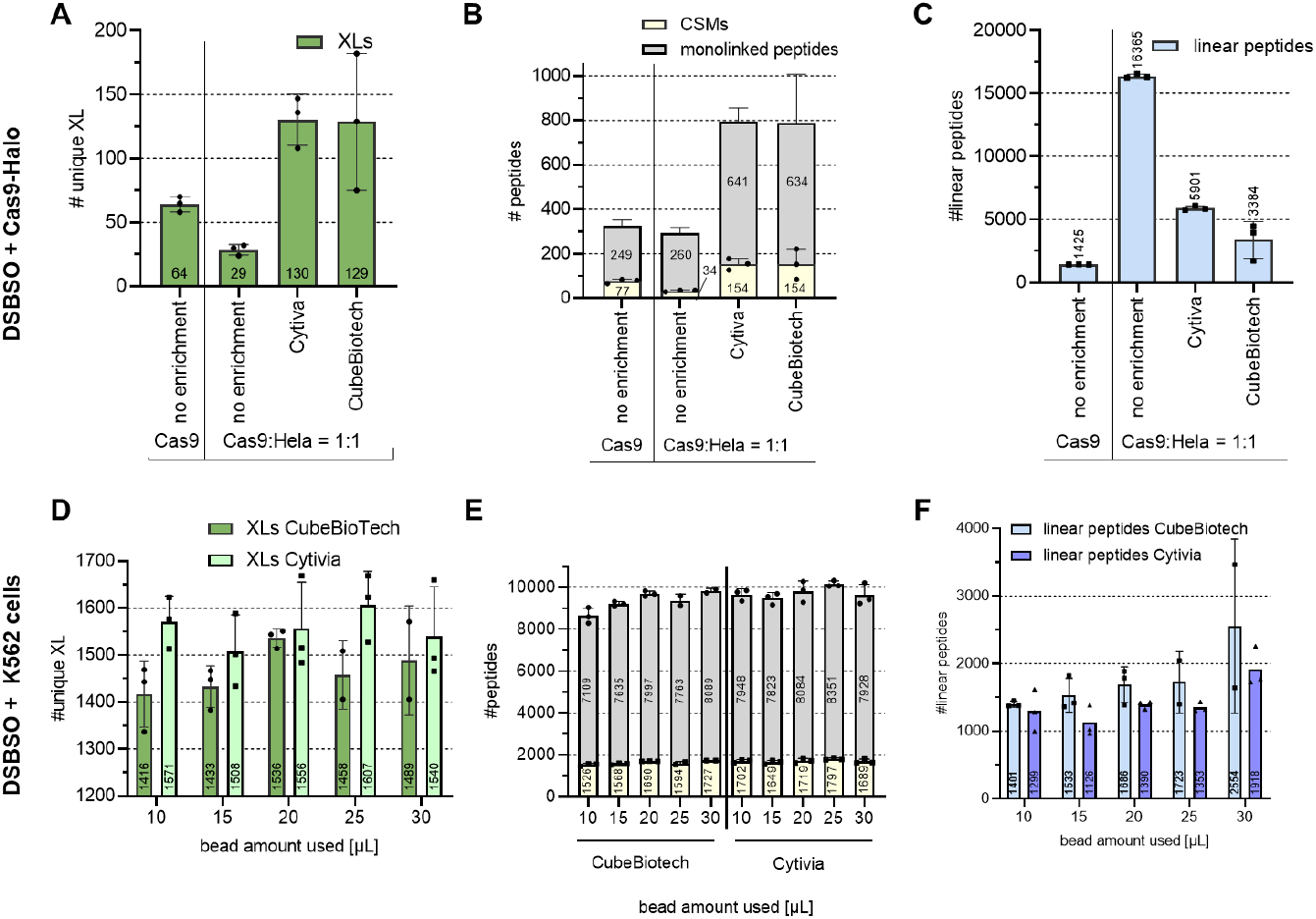
Benchmarking magnetic beads from different vendors & finding optimal enrichment conditions: **A-C**: 20 µg DSBSO crosslinked Cas9-Halo peptides, spiked into a non-crosslinked background from human HeLa peptides (1:1, w:w) were either enriched using magnetic beads from Cytivia or Cube Biotech as indicated using 20µL bead slurry and using the exact same processing workflow. Data was searched against a database containing 20585 proteins (Cas9 + human proteome). **D-F**: Living K562 cells were crosslinked and enrichment was applied using both bead types under variation of the used bead slurry volume as indicated. Data was searched against the human proteome. Bars indicate identified average numbers of XLs (**A, D**), CSMs & monolinked peptides (**B, E**) or linear peptides (**C, F**) at 1% FDR level, and error bars indicate their standard deviation, n=3

Next, we extended our tests to *in vivo* crosslinked K562 cell samples corresponding to 0.25 mg protein content. We titrated the amount of bead slurry used for enrichment and found a surprisingly small correlation of applied bead amount on resulting CSMs/ unique crosslink sites (Figure 1D-E). Additionally, only a minor increase of non-specific binders upon increasing the excess of bead material (Figure 1F) was observed. Given the better properties concerning handling, slightly improved XL numbers and slightly reduced background of linear peptides, we chose to proceed with the Cytivia beads for all subsequent studies. We identified 25 µL bead volume as optimal condition between having enough binding groups and offering a higher surface for unspecific binding. Of note, this correlates to a bead volume/ protein ratio of 100µL/mg protein and this ratio was also kept for the subsequent upscaled studies.

### Proof of principle study in K562 cells

We applied the now refined affinity enrichment settings to the full workflow for *in vivo* DSBSO crosslinking adopted from the work of the Huang lab^9^ (b). To streamline the workflow and potentially reduce sample losses on the (long) way from the cell culture to the mass analyzer we aimed to further streamline the workflow by reducing the number of processing steps. Omitting C18 cleanup ahead of the affinity enrichment step led to only slightly reduced identifications of crosslinks by 13 % (Figure 2 A-B) while unexpectedly helped to reduce the background from linear peptides by ∼38% on average (Figure 2C). We hypothesize, that the presence of 0.5M NaCl (added in elution buffer of FASP) did not adversely affect the robustness of the click reaction, while it lowered unspecific binding of peptides to the bead-surface. When skipping the C18 cleanup step and applying size exclusion chromatography (SEC) as an orthogonal enrichment strategy instead, overall workflow sensitivity was dramatically improved leading to >3000 unique crosslinked sites identified on average from K562 cells or nuclear extracts from K562 cells and within 4 SEC fractions measured on the LC-MS (Figure 2D). This improved sensitivity seems to mainly originate from reduced sample complexity with a lower background of linear peptides (Figure 2F). In contrast, the obtained average ratio of CSMs/ monolinked peptides initially appears to remain at ∼0.2 -0.25, independent of SEC enrichment (Figure 2E and 2B). However, a sufficient separation of monolinked from crosslinked peptides was facilitated via SEC specifically in the early fractions, containing more and larger crosslinked peptides. In these early fractions the CSM/monolink ratio was improved to >6 and drops to <0.2 for later eluting fractions (Supplemental Figure 2). Of note, a complete separation of monolinked from crosslinked peptides was not expected and our results successfully reproduce previous reports for SEC based crosslink enrichment.^12^ In this context, we noticed that the vast majority of crosslink IDs, namely ∼87% & ∼90% when crosslinking the entire cell or the enriched nuclei, respectively, were found in only one SEC fraction (Figure 3A-B). In conclusion, although fractionation was performed in this workflow, we propose that only a single fraction suffices for comparable coverage. As MS acquisition time is often limited and expensive, this strategy demonstrates another way to streamline this workflow especially for larger sample cohorts in biological/medical studies.

**Figure 2:**
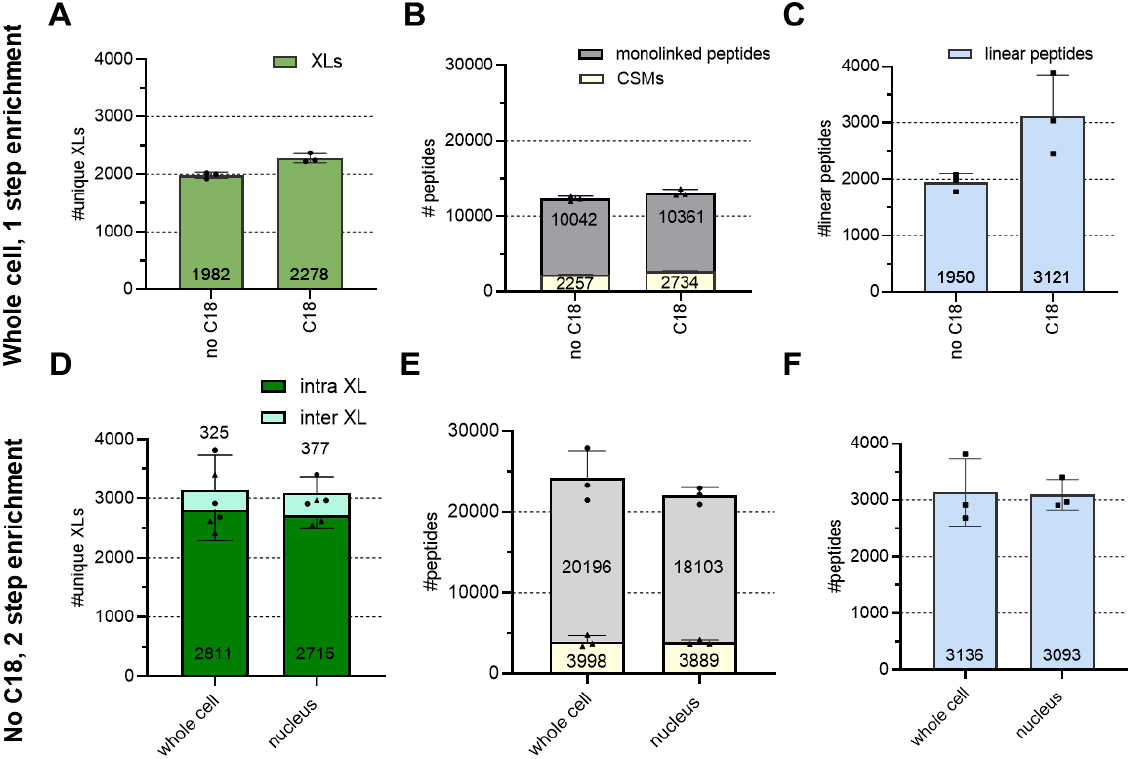
Proof of principle study – DSBSO in vivo: **A-C**: Living K562 cells were DSBSO crosslinked and processed with or without a C18 cleanup step ahead of affinity enrichment using the Cytivia beads but without an additional enrichment. **D-F**: Whole K562 cells or their nuclear extracts were DSBSO crosslinked employing a streamlined workflow without C18 enrichment but including orthogonal SEC fractionation. Bars indicate identified average numbers of XLs after combined analysis of 4 SEC fractions (**A, D**), CSMs & monolinked peptides (**B, E**) or linear peptides (**C, F**) at 1% FDR level, and error bars indicate their standard deviation, n=3 independent replicates

**Figure 3:**
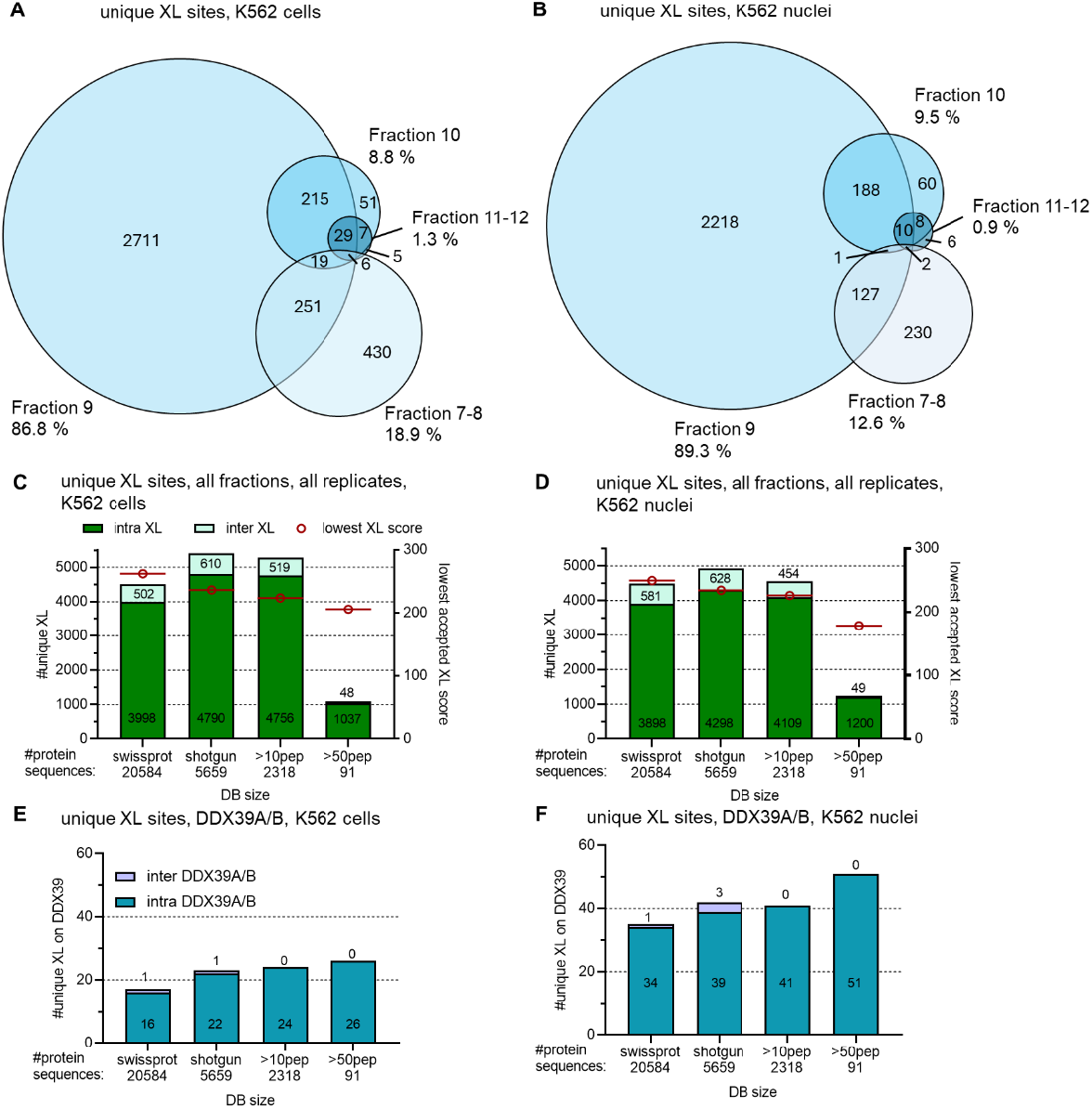
Improving coverage with minimal effort. Venn diagrams show commonly identified unique XL sites across the acquired SEC fractions from a representative replicate each from whole K562 cells (**A**) and K562 nuclei (**B**) respectively. **C-F**: Changes of crosslink ID numbers depending on database size. Bar plots showing either all XL unique XL sites from whole cells (**C**) and nucleus (**D**) or within those only XL unique XL sites found on either DDX39A or B from whole cell (**E**) or nuclear extract (**F**) samples. All fractions and n=3 replicates for each condition searched together at 1% FDR.

To achieve maximal coverage of crosslink matches from our data, we decided to run a shared analysis of files from all fractions towards databases of different sizes (Figure 3 C, D). Thus, we first generated a shotgun database from n=5 biological replicates of non-crosslinked K562 samples, yielding 5659 proteins. From that we created smaller databases containing only sequences from proteins of which at least 10 or 50 peptides were found in the shotgun data, yielding 2318 and 91 remaining proteins, respectively. 19 and 20 unique peptides were found from the main proteins of interest in this study, DDX39A and its homologue DDX39B, but both proteins were kept in the database even for the most stringently filtered database requiring 50 peptides/protein. With this we aimed to maximize crosslinks potentially detectable on those two proteins. As shown in Figures 3 C and D, using the full shotgun database was most advantageous in both systems, whole cells and nuclear extracts, to maximize overall crosslink numbers. We hypothesize, that this is because of less possible random (decoy) hits when using a reduced database size, resulting in a less stringent score cutoff applied (dropping by ∼21% and 28 % for whole cell and nucleus samples respectively, see Figure 3C, D) while maintaining a 1% FDR threshold. For DDX39, reducing the database size led to slightly higher intra-link numbers for DDX39 proteins (Figure 3 E, F) while lowering overall numbers, likely due to crosslinked proteins being removed from the database. No additional inter-protein hits for DDX39 were found using databases smaller than the 5659-protein sized shotgun database. Simultaneously the risk of less confident intra-links being reported as false positives is higher with very small databases. Consequently, we decided to continue with data from the full shotgun database for further data processing.

### Study of DDX39B

The ATPase DDX39 is a central molecular switch that directs mature mRNA ribonucleoprotein complexes and thereby controls mRNA export.^13^ To validate the expected primary localization of DDX39B in the nucleus within the herein used cell-based system we made applied a clone expressing the GFP tagged protein version. DNA was counterstained using DAPI confirming co-localization with DDX39B as expected (Figure 4 A). In conclusion we decided to focus on samples measured from nuclear extracts where higher concentrations of DDX39 are expected (as confirmed by yielding higher XL-ID numbers, shown in Figure 3 C vs D).

**Figure 4:**
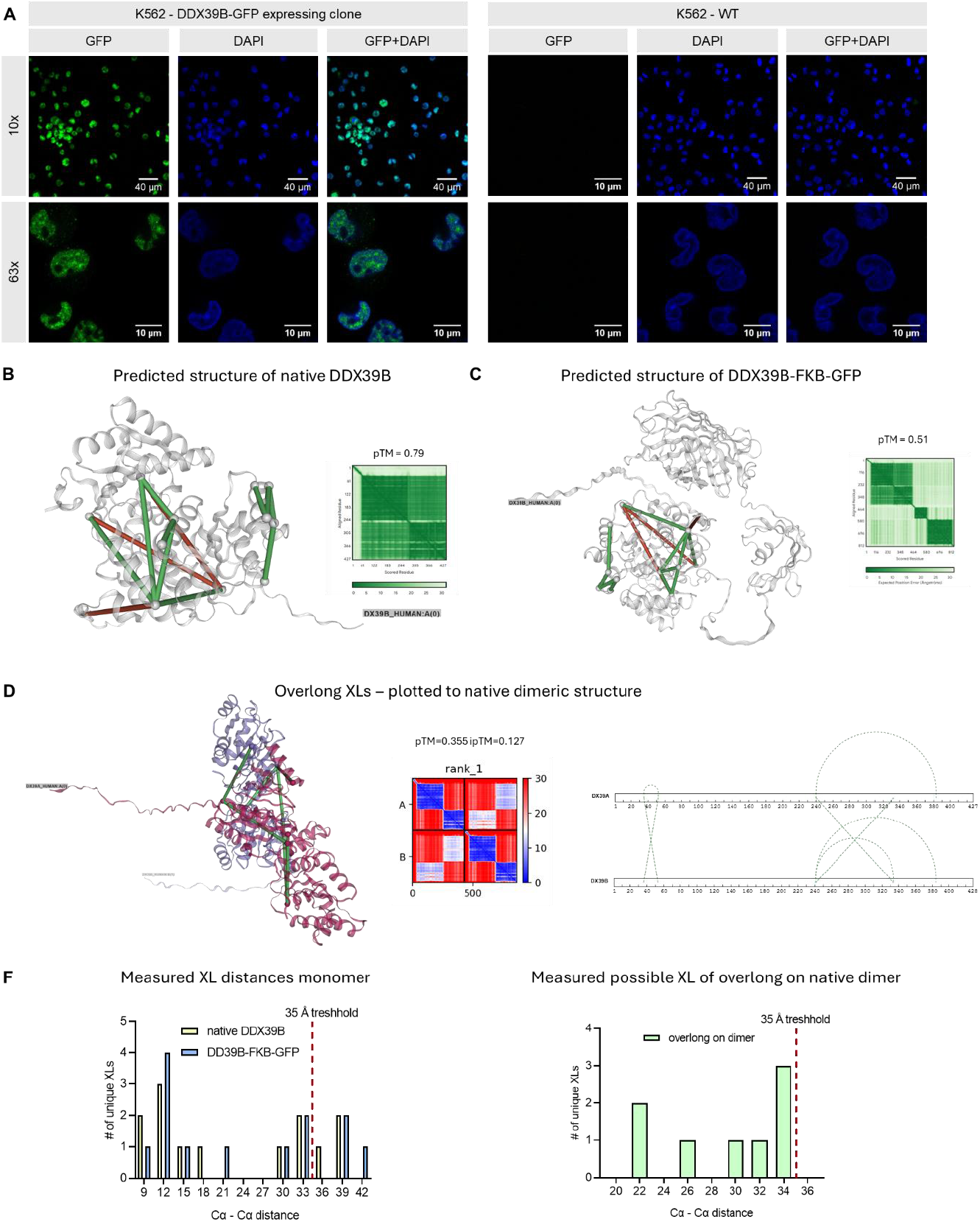
Localization & structure prediction of DDX39. **A:** DDX39B-GFP expressing and wt-K562 cells were DNA stained using DAPI and inspected by fluorescence microscopy at 10x or 63x magnification as indicated. GFP signal shown in green and DAPI signal shown in blue. **B, C**: Crosslinks found within the nuclear extracts/ analyzed using the shotgun database plotted on the native DDX39B (**B**) or the tagged DDX39B-FKB-GFP (**C**) rank 1 3D structure model predicted from AlphaFold3 and their respective diagnostic plots. For ambiguous crosslinks only the shortest possible connection, when plotting to the predicted 3D structure from AlphaFold2 is shown. Shown in green are crosslinks ≤35 Å and in red links >35 Å. **D**: Possible K-K connections with ≤35 Å from previously non satisfied crosslinks after plotting onto the monomeric DDX39B structure, now plotted to the predicted structure of the DDX39A & B complex and allowing links to fit on either protein or across both proteins. **F**: Histogram of measured Cα to Cα distances when plotting found crosslinks to the predicted 3D structures as indicated.

Of those 39 identified intra-protein crosslinks for DDX39A or B, most could not unambiguously be annotated to only one homologous protein sequence (see Figure 5 A, with unambiguous links shown as dashed line). We therefore first focused on DDX39B and predicted its structure with or without the attached FKB-GFP tag using AlphaFold 3^14^. To remove ambiguous hits, we further selected only the shortest possible crosslink distance as measured on this predicted structure for plotting and further processing (Figure 4 B-C). We decided on applying a distance cutoff of 35 Å as suggested by Jiao et al.^9^ leading to ∼77 % of unique DSBSO links in agreement with the predicted structure. The number of overlong crosslinks was not altered in dependence of the FKB-GFP tag included for prediction. Of note, the predicted template modelling (pTM) score is lowered for the tagged prediction vs the native from 0.79 to 0.51 respectively and the resulting average crosslink length was slightly elongated in presence of the tag (Figure 4F). All distance measurements were performed using the respective rank 1 models, with only medium to low pTM scores though as described above. (Figure 4). Notably, since DDX39A and DDX39B are highly homologous, all overlength crosslinks found are ambiguous and might relate to an intra-protein connection on DDX39A or to an inter-protein connection across DDX39A and B in dimeric form. We predicted the structure of the respective dimer, yielding a dimeric structure with relatively low pTM and iPTM scores (Figure 4D), which might be reasoned by additional biologically relevant ligands such as RNA not considered for prediction. However, our experimental data suggests, that the predicted dimeric structure could be indeed valid with crosslink connections satisfying the maximum distance threshold for DSBSO for all previously overlong links (Figure 4D). We therefore assume a monomeric and dimeric protein form present within the nuclear extracts both captured in our data. Of note, our findings are in line with previous reports describing that overexpressed DDX39A can heterodimerize with DDX39B which would inhibit its function.^15^

**Figure 5:**
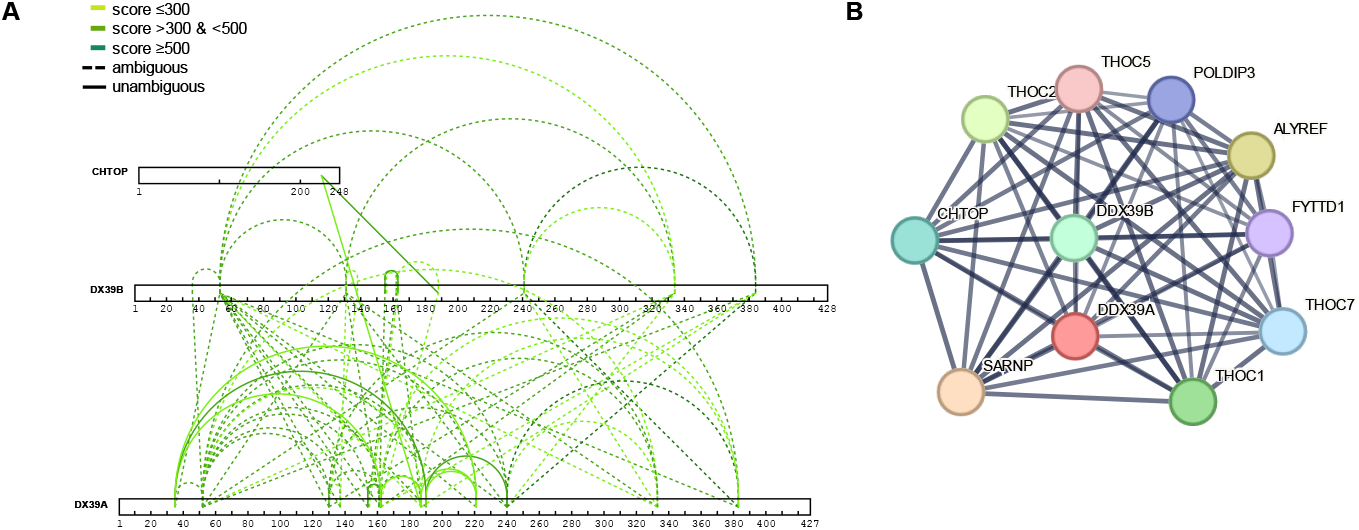
Identified PPI of DDX39 A/B. **A**: All XLs found on DDX39B (shown without tag) and its direct interaction partners. Dashed lines indicate ambiguous links which might originate from intra-protein links or inter-protein links to the homologous sequence of DDX39A/B respectively. Solid lines indicate unambiguous XLs and darker green indicates higher confidence. **B**: Known interaction partners of the DDX39A/B complex as given in the STRING database.

In addition to these DDX39 intra-complex connections, we detected crosslinks to CHTOP (Figure 5 A). We compared our findings to known PPI for DDX39 using the STRING database^16^ (Figure 5 B) and found that a physical interaction of DDX39 A and B and its interaction to CHTOP are known, which we confirmed in this study. Our data demonstrates an interaction of both DDX39A and B to K226 of CHTOP which confirms earlier results by Fasci et al.^17^, who identified this exact crosslink connection in nuclear extracts from U2OS cells using a different crosslinker (DSSO).

### A comprehensive PPI network

In addition to the data generated on DDX39 we obtained a comprehensive PPI network from the used K562 model system. A detailed list of all crosslink matches can be found in Supplemental Tables 1 & 2 for results searched towards the shotgun database from whole cells and nuclear extracts respectively. Identified crosslinked proteins were distributed across several cellular compartments with the vast majority found within the cytosol or nucleus (Figure 6A). When crosslinking within nuclear extracts, ∼80% of all crosslink matches could be annotated to the nucleus, which enables purity estimation of our samples after nuclei enrichment. Reported proteins present within the nucleus or the cytosol according to their GO-annotations are depicted as such in Figure 6A And were counted as co-localization. As little as 0.5% and 0.2% for whole cell and nuclear extract respectively, of all crosslinks were between proteins from different cellular compartments. Such crosslinks theoretically should not form. Although of course no proof for FDR, the prevalence of non-co-localizing crosslinks way below 1 % indicates a potentially functional FDR control, while a higher prevalence would be concerning. Within the whole cell dataset, analyzed towards a generated shotgun database (see Figure 3C), we found a total of 311 PPI from non-ambiguous crosslinks including networks within the ribosomal complexes, chaperonin-containing T complex, 48S & 43S preinitiation complexes and others (Supplemental Figure 4, Supplemental Table 4) Within the nuclear extract dataset we found a total of 393 PPI from non-ambiguous crosslinks including networks within the ribosome; spliceosome, transcriptional export proteins and others (Figure 6B). Our data thereby provides valuable insights into molecular pathways within the cell and includes 56 (14% of all PPI) potential novel interaction partners, not listed in the String database (Supplemental Table 3). In line with the findings suggested by Fürsch et al^18^, our data reports crosslinks primary on more abundant proteins within the cell (Supplemental Figure 3). However, our crosslink-data covers ∼5-6 orders of magnitude in abundance of crosslinked proteins showcasing that at least some crosslinks even from low abundant proteins have been pulled out thanks to the effective enrichment strategy. Moreover, within the nuclear extract we were able to go cover more low abundant proteins compared to the whole cell crosslinking approach (Supplemental Figure 3).

**Figure 6:**
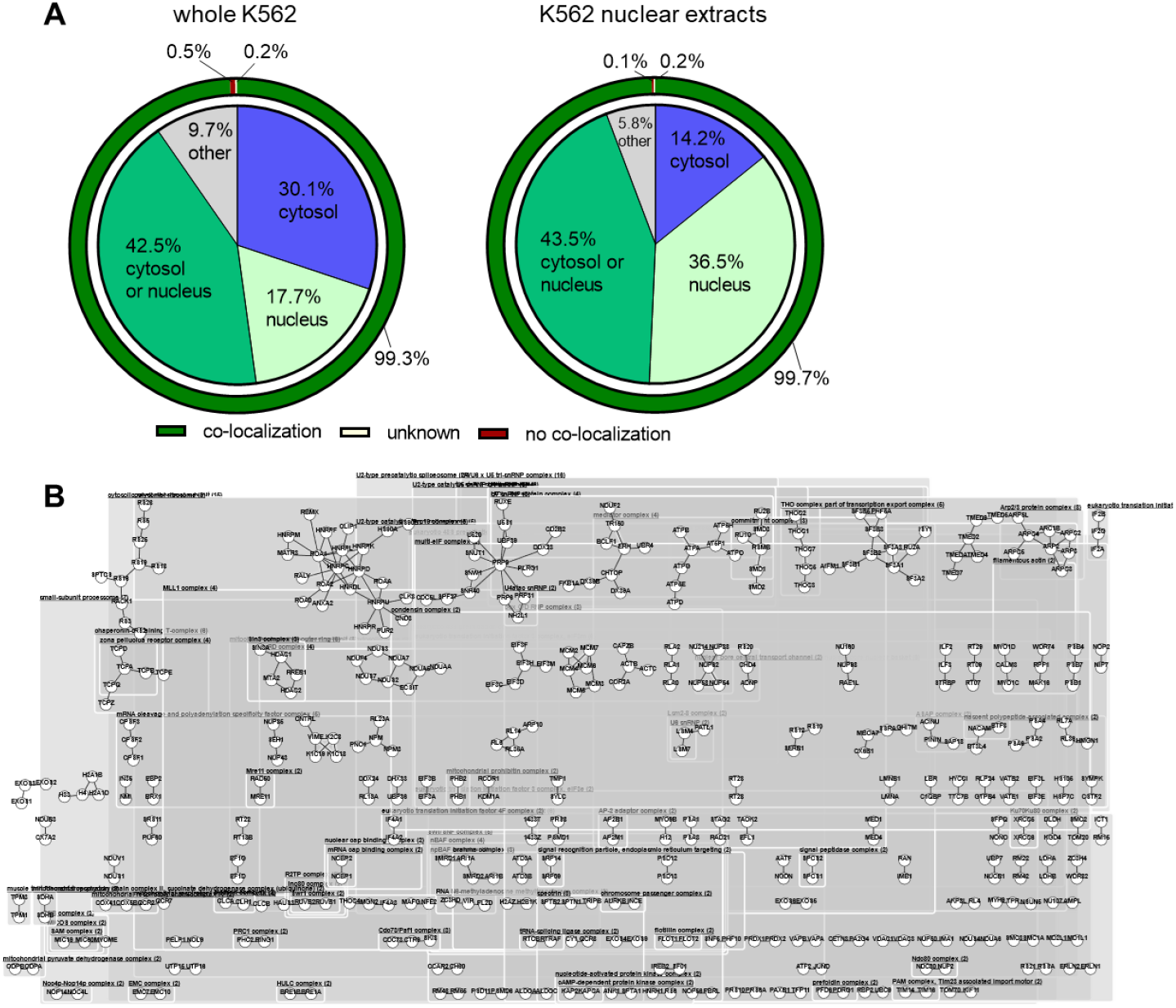
PPI found from in vivo XL within K562 cells. **A**: Distribution of cellular compartments of crosslinked proteins after crosslinking intact cells or their nuclear extracts as indicated. Data based on go-annotations of DSBSO connected proteins. The outer sphere shows the fraction of connected proteins within the same cellular compartment (co-localization) or within different compartments. **B**: PPI network as found from nuclear extract crosslinking samples after analysis against a shotgun database (see Figure 3C) with correlated groups annotated. In total 333 non-ambiguous PPI from 628 heteromeric crosslinks are shown.

## METHODS

Figure 7 provides an overview of the workflow, with detailed explanations given in the following sections.

**Figure 7:**
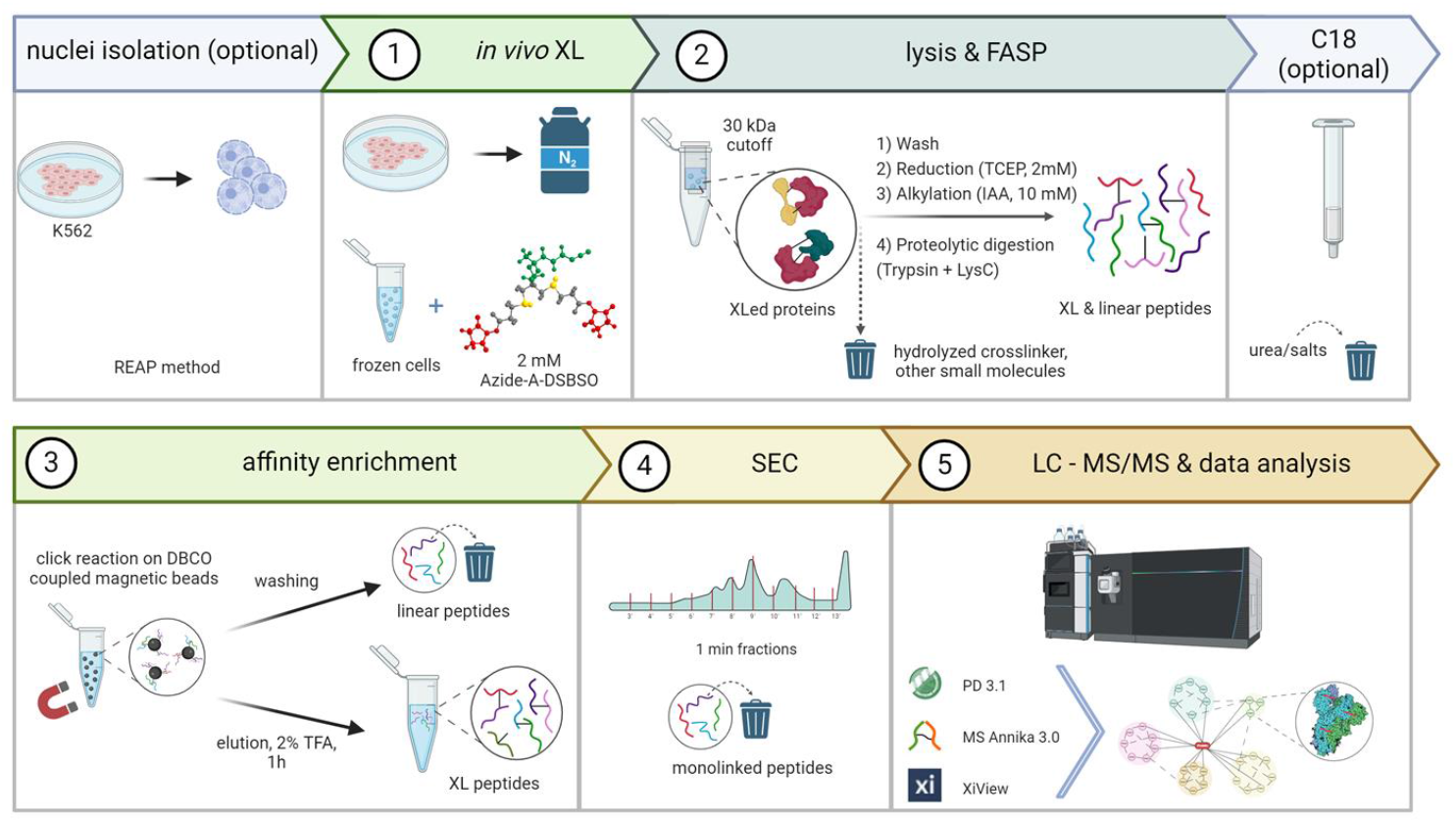
Graphical workflow representation.

### Reagents

Purified recombinant Cas9 from *S. pyogenes* fused with a Halo-tag was generated in house, as described by Deng et al.^19^ Azide-A-DSBSO was obtained from Sigma-Aldrich (#909629). For enrichment bead generation, dibenzocyclooctyne-amine (DBCO-amine, #761540, Sigma-Aldrich) was coupled to NHS Act Sepharose® 4 Fast Flow (Cytivia, #GE17-0906-01), NHS Mag Sepharose® (Cytivia, #28951380) beads, or Pure Cube NHS act. MagBeads (Cube Biotech, #50405) as indicated. The prepared beads were stored as 50% slurry in a 1:1 ethanol:water mixture. Trypsin gold was purchased from Promega (Mannheim, Germany) and lysyl endopeptidase (LysC) was from Wako (Neuss, Germany). Benzonase—pharmaceutical production purity—was purchased from Merck (Darmstadt, Germany). All other solvents were purchased from Fisher Chemicals if not indicated otherwise.

### Cell Culture and Nucleus Preparation

K562 wild-type (WT) and DDX39B-dTag-GFP mutant cells were grown in RPMI 1640 medium, supplemented with 10% fetal bovine serum (FBS), 1 mM sodium pyruvate, 1% penicillin/streptomycin, and 2% L-glutamine. The cells were cultured at 37°C in a humidified incubator with 5% CO_2_. Once the cultures reached the appropriate density, the cells were washed with phosphate-buffered saline (PBS), divided into aliquots, and stored in liquid nitrogen for future use.

To isolate nuclei, a modified Rapid, Efficient, and Practical (REAP) protocol was used. The cells were lysed gently in ice-cold PBS containing 0.1% Igepal CA-630. Nuclei were collected through centrifugation and washed to remove any remaining cytoplasmic components. The purified nuclei were aliquoted and stored at −80°C for further experiments.

### XL reaction Cas9

Purified recombinant Cas9 from S. pyogenes fused with a Halo-tag at 1 mg/mL in 200 mM HEPES (pH 7.6) was crosslinked by adding a 4mM DSBSO in dry DMSO stock to a final concentration of 0.2 mM DSBSO. The crosslinking reaction was incubated at 37°C on a thermoshaker set to 1000 rpm for 1 hour. Crosslinked Cas9 samples for initial check-experiments were either used without additional enrichment or spiked in 1:1 (w/w) ratio into non crosslinked Pierce™ HeLa Protein Digest (Thermo Fisher) followed by DBCO bead-based enrichment as indicated.

### Chemical Crosslinking of K562 Cells and Nuclei

Previously frozen cells or nuclei were thawed quickly and resuspended in a crosslinking buffer consisting of 20 mM HEPES (pH 7.6), 150 mM NaCl, and 1.5 mM MgCl_2_. Protein concentrations were standardized to approximately 13 mg/mL for whole cells and 5 mg/mL for nuclei. Crosslinking was initiated by directly adding the DSBSO powder to the cells/nuclear extracts to reach a final concentration of 2 mM DSBSO. Samples were incubated shaking at 37°C for 1 hour at 1000 rpm. 1µL of Benzonase was added for the final 30 minutes during the crosslinking reaction to facilitate DNA digestion. No quenching of crosslinking was performed as potential excess & reactive linker was removed by FASP immediately thereafter.

### Protein Quantification and Filter-Aided Sample Preparation (FASP)

The samples were lysed using a Bioruptor sonicator set to 5-minute cycles (30 seconds on and 30 seconds off) at 4°C. After lysis, the samples were clarified by centrifugation at 14,000 × g for 15 minutes at 4°C and kept on ice until further processing.

Protein concentrations were measured using the Bio-Rad Protein Assay Dye Reagent (Cat. No. 5000006). Each replicate required 1 mg of protein, which was divided into three FASP filters (0.33 mg per filter; Merck Millipore, Cat. No. MRCF0R030). The peptides eluted from these filters were pooled to obtain 1 mg of peptides per replicate.

Filters were equilibrated with 8 M urea in 0.1 M Tris-HCl (pH 8.2) by centrifugation at 14,000 × g for 15 minutes. Proteins were loaded onto the filters in the same buffer and concentrated by centrifugation, followed by a wash step with equilibration buffer. Reduction was carried out using 2 mM Tris(2-carboxyethyl)phosphine (TCEP) in 8 M urea, and alkylation was performed with 10 mM iodoacetamide (IAA) in 8 M urea, both at room temperature. After this step, the workflow diverged: For the general FASP approach, filters were washed twice with 100 mM HEPES (pH 7.3) to remove residual reagents, while for the modified FASP with C18 clean-up, an alternative washing protocol was applied, as described in the respective section.

Proteins were digested sequentially, starting with lysyl endopeptidase (LysC) at a 1:100 (w/w) enzyme-to-protein ratio in 100 mM HEPES (pH 7.3) at 37°C overnight, followed by trypsin digestion under identical conditions for 4 hours. Peptides were first eluted by centrifugation, followed by sequential elution with 100 mM HEPES (pH 7.3) and 0.5 M NaCl. The eluates from the three filters were pooled to generate a final peptide sample.

### Modified FASP with C18 Clean-Up

For the modified protocol, filters were first washed with 25 mM ammonium bicarbonate (ABC) buffer (pH 9.2) containing 8 M urea before digestion. LysC digestion was conducted in 5.45 M urea and 25 mM ABC buffer (pH 9.2), while trypsin digestion was performed in 1.5 M urea and 25 mM ABC buffer. The peptides were eluted as described and then further purified using Sep-Pak® tC18 cartridges (Waters, Cat. No. WAT054960).

Sep-Pak® tC18 cartridges were equilibrated with a sequence of methanol, 70% (v/v) acetonitrile (ACN) with 0.1% trifluoroacetic acid (TFA), and 0.1% (v/v) TFA. Acidified samples (adjusted to pH 3–4 with TFA) were loaded onto the cartridges. After washing with 0.1% (v/v) TFA, peptides were eluted using 70% (v/v) ACN containing 0.1% (v/v) TFA. To prevent DSBSO acetal hydrolysis, the eluates were neutralized with 1 M HEPES (pH 7.3). Finally, the samples were concentrated using a SpeedVac and reconstituted in 100 mM HEPES (pH 7.3) to ensure consistency with the conditions used in the standard FASP procedure.

### Preparation of DBCO coupled beads for affinity enrichment

The NHS Mag Sepharose® beads manufactured by Cytivia were used for all *in vivo* XL studies. 500 µL of the bead slurry (∼20% slurry) were washed immediately prior to usage once with 1 mL ice-cold 1mM HCl. After that, beads were resuspended in a mixture of 385 µL 50 mM HEPES pH 7.5 and a stock solution of 64 µL 10 mg/mL Dibenzocyclooctyne-amine (761540-10MG, Sigma) in dry DMSO. The resulting suspension was incubated for 2 hours at RT on a rotator followed by 2x washing using 50 mM HEPES pH 7.5, 1x washing with 1M NaCl and 2x washing with 100 mM Tris-Cl pH 7.5. For storage at 4°C, beads were resuspended in 100 mM Tris pH 7.5 to reach a final volume of 1000 µL.

### Affinity enrichment of XL-peptides

Beads for affinity enrichment were extensively washed to ensure optimal binding conditions. Initially, they were washed five times with 1 M NaCl in 50 mM HEPES (pH 7.3) to remove contaminants, followed by five additional washes with 50 mM HEPES (pH 7.3) for equilibration.

Samples were incubated with prepared beads under rotation for 2 hours at room temperature or overnight at 4°C, depending on the experiment. For bead optimization, ratios of 10, 15, 20, 25, and 30 μL of beads per 0.25 mg of protein were tested, with 25 μL per 0.25 mg found to be optimal. This condition was later upscaled (4x) to 100 μL of beads per 1 mg of protein for all *in vivo* studies. Cas9 tests were performed with 20 μL of beads per 20 μg of recombinant Cas9 protein, with an additional 20 μg of HeLa digest for spiking experiments, mimicking cellular background peptides.

After incubation, the supernatant was removed using magnetic separation. Beads were washed sequentially with a volume five times greater than the bead volume. The washing protocol included five washes with 1 M NaCl in 50 mM HEPES (pH 7.3), followed by three washes with 10% (v/v) acetonitrile (ACN), and four washes with MilliQ H_2_O (0.2 µm filtered). To ensure thorough cleaning, the beads were transferred to a new tube for a final wash with MilliQ H_2_O, minimizing any potential carryover before elution.

Crosslinked peptides were eluted in two steps using 2% (v/v) trifluoroacetic acid (TFA) in water. In the first step, two-thirds of the bead slurry volume was incubated for 1 hour, followed by elution of the remaining one-third for 30 minutes. The eluates were combined, and those intended for immediate analysis were dried using a SpeedVac and resuspended in 0.015% DDM in 0.1% TFA. These samples were either resolved at 4°C overnight or at room temperature for at least 30 minutes prior to analysis. For samples designated for size exclusion chromatography (SEC), the final concentration step adjusted the sample volume to ∼45 μL to ensure compatibility with SEC workflows.

### Fractionation via SEC

SEC was performed using a TSKgel SuperSW2000 column (4 µm particle size, 4.6 mm ID × 30 cm) to separate peptides based on size. The column operated at a flow rate of 0.3 mL/min with detection at 214 nm. The elution buffer consisted of isocratic elution with 30% (v/v) ACN and 70% (v/v) water, both containing 0.1% (v/v) TFA. Samples were injected in their entirety, and fractions were collected at 1-minute intervals.

Fractions were pooled based on the intensity of the elution peaks. Fractions with lower peptide intensity were combined (e.g., 7+8 and 11+12), while those with higher intensity (e.g., fractions 9 and 10) were processed separately. Prior to MS/MS analysis, all samples were resolved in 0.015% DDM and 0.1% TFA and injected at a final volume of 10 µL.

### Liquid Chromatography–Mass Spectrometry (LC–MS) Analysis

Analyses were performed using a Vanquish Neo UHPLC system (Thermo Fisher Scientific). Samples were loaded onto a PepMap Acclaim C18 trap column (5 mm × 300 μm, 5 μm particle size, 100 Å pore size; Thermo Fisher Scientific) in trap-and-elute mode, then switched in-line with the analytical column for separation. For bead benchmark studies, separation was carried out on a PepMap Acclaim C18 analytical column (500 mm × 75 μm, 2 μm, 100 Å; Thermo Fisher Scientific) at a flow rate of 0.300 μL/min. All other analyses were conducted on an Aurora Ultimate 25 cm column (250 mm × 75 μm, 1.9 μm, 120 Å; IonOptiks) at 0.230 μL/min. The column temperature was maintained at 45°C.

A linear gradient was applied, gradually increasing Buffer B from 2% to 45% over 120 minutes, followed by a transition to 95% Buffer B within a minute. The column was maintained at 95% Buffer B for 6 minutes before re-equilibration at 2% Buffer B. Buffer A consisted of 100% water with 0.1% (v/v) formic acid, while Buffer B was composed of 80% (v/v) acetonitrile with 0.1% (v/v) formic acid.

Mass spectrometric analyses were performed on an Orbitrap Eclipse Tribrid mass spectrometer (Thermo Fisher Scientific) equipped with a FAIMS Pro interface set to standard resolution, with compensation voltages of –50 ± 10 V. The instrument was operated in positive-ion mode under a data-dependent acquisition (DDA) Top 10 method. MS1 scans were acquired over an m/z range of 375–1500 at a resolution setting of 120,000, with an automatic gain control (AGC) target of 1.2 × 10^6^ (300% normalized). MS2 scans were generated using HCD at stepped collision energies of approximately 27 ± 6%, with a 1.2 m/z isolation window, a resolution of 30,000, and an AGC target of 5.0 × 10^4^ (100% normalized). Precursor ions were restricted to charge states 3–8, required a minimum intensity threshold of 2.5 × 10^4^, and were dynamically excluded for 30 minutes after one fragmentation event. The spray voltage was set to 2.3 kV, and the capillary temperature was maintained at 275°C.

### Fluorescence Microscopy

All imaging was performed on a Zeiss LSM 780 confocal microscope in laser scanning confocal mode. High-magnification images for nuclear colocalization were acquired using a Plan-Apochromat 63×/1.40 Oil DIC objective, while overview images were taken with a Plan-Apochromat 10×/0.3 M27 objective. EGFP (DDX39B-GFP) fluorescence was excited at 488 nm and detected at approximately 517 nm, and DAPI was excited at 405 nm with emission collected near 451 nm. The pinhole was set to roughly 1 Airy unit for both channels, and laser power attenuation ranged from about 96 to 99 %. Detector gains were typically set between 822–852 for EGFP and 707–757 for DAPI, with digital gains of 3.86× (EGFP) and 3.19× (DAPI). Z-stacks of seven slices were acquired per field of view to capture nuclear volumes. Image processing was done via ImageJ Version 1.54m.

### Data Analysis

LC-MS/MS raw files were analyzed within Proteome Discoverer 3.1 (Thermo Fisher Scientific), using MS Amanda 3.0 ^20^ for searching linear peptides and MS Annika 3.0^21^ to search crosslinked peptides. Relative, label free, quantification was done using apQuant^22^. Results were filtered to 1% FDR at peptide, protein, CSM and unique XL site level. A DSBSO linker based workflow as recommended in the most recent MS Annika publication^21^ (download: https://github.com/hgb-bin-proteomics/MSAnnika/raw/master/workflows/PD3.0/DSBSO_MS2.pdAnalysis) was used for data analysis. Searches were performed towards databases of different sizes as indicated for each result within this study individually.

### Post Processing

For data visualization and statistics Graph Pad Prism 8.0 (GraphPad Software Inc.) was used. Venn diagrams were plotted using DeepVenn^23^. Post processing to visualize interaction networks or 3D models was performed using xiVIEW^24^. Results obtained from Annika in Proteome discoverer were exported to xiview using a custom Python script available on GitHub (https://github.com/hgb-bin-proteomics/MSAnnika_exporters/blob/develop/xiViewExporter_msannika.py). 3D structure prediction was performed using AlphaFold3^14^ on a local cluster. AlphaFold2^25^ on a Google Colab sheet^26^ was used to predict the dimeric DDX39A-B structure using the suggested default settings (https://colab.research.google.com/github/sokrypton/ColabFold/blob/main/AlphaFold2.ipynb?pli=1#scrollTo=R_AH6JSXaeb2).

## DISCUSSION

Despite tremendous improvements already made in the field of crosslinking mass spectrometry, with a multitude of different linker chemistries and sample preparation protocols being developed, the generation of comprehensive *in vivo* crosslinking data is still challenging. Within this study, we provide an easy to use, streamlined workflow for this purpose. We provide guideline of which magnetic bead type is best suited, and at which excess over DSBSO crosslinker enrichment beads perform optimal. By removing a C18 based purification step we reduce potential sample losses without suffering from relevant final crosslink coverage. This is especially advantageous for studies with limited sample input. While applying an orthogonal SEC based enrichment helped to boost crosslink IDs to an overall combined number of >5000 unique XLs on 1692 proteins in the whole cell samples. Its tricky to compare yielded coverage to other studies as conditions such as cell type, linker type, sample preparation workflow, LC-MS settings, data analysis software and search settings differ a lot. However, to give an example from another comprehensive study in K562 cells, Yugandhar et al^27^ found 9300 crosslinked residues of which 1268 were interprotein crosslinks, using DSSO. While this outperforms the numbers seen in this study, they performed extensive SCX fractionation and generated 122 raw files to obtain such an impressive result. In contrast only 12 raw files yield to the combined result in this study and of those 5000 unique XLs found, more than 3000 unique XLs on average were found per replicate (in 4 raw files). Our data further shows that even without fractionation and using only a single step affinity-enrichment (hence suffering from least potential sample losses) around 2000 unique crosslinked sites from *in vivo* crosslinked K562 cells are yielded. Our data further shows that also after SEC fractionation, measurement of only one fraction is sufficient to yield close to 90 % of all links found in all fractions which is still a substantial boost of crosslink coverage by around 27% to > 2500 unique crosslinks, compared to the 1 step enrichment, with the additional benefit of saved MS measurement time.

We generated a comprehensive proteome-wide PPI network from K562 cells as well as a network from their nuclear extracts, containing 56 (14 % of total) novel PPIs not listed in the STRING database. Our data can therefore be used to screen for novel interactors of interest or as a complementary dataset confirming orthogonal biochemical approaches. We additionally present that performing the crosslinking reaction within a cellular sub-compartment (i.e. nucleus) helps to improve finding crosslinks on proteins not abundant enough otherwise.

Of note, our attempts to improve predicted protein structures of DDX39 by integrating crosslink data using AlphaLink2 yielded in altered but not necessarily improved protein-complex structures with lowered ipTM scores and an increased fraction of dissatisfied crosslinks. We hypothesize that the high homology of both protein sequences (DDX39A and B) in automatic sequence alignment yielded suboptimal results. With an altered sequence not identical to the original sequence of DDX39B, predicted structures potentially did not fit equally well to the obtained crosslink data. Furthermore, multiple protein conformations present in solution might make it impossible to find a single protein complex structure satisfying all false positive crosslinks at a time.

The here presented easy to implement comprehensive in vivo crosslink workflow will be highly valuable for the community including non-crosslink-experts, who can apply this simple workflow to any cell type or enriched subcellular fraction with general wet lab skills. The identified shared PPI network displayed in this study can be further used as a general resource for data mining and will serve as complementary dataset for (novel) PPIs identified with orthogonal approaches in the future.

## ACKNOWLEDGMENTS

The authors would like to thank Julius Brennecke and Nuno Maulide for their support and fruitful discussions throughout this project. Furthermore, we would like to express our gratitude to Micha Birklbauer for his continuous support using Annika, and his help to resolve export and data processing problems. This work was supported by the infrastructure funding 4th call 2022/01 (AT-SCP) of the Austrian Research Promotion Agency (FFG) and the project LS20-079 of the Vienna Science and Technology Fund (WWTF). This work was further funded by the P35045-B project (Grant-DOI 10.55776/P35045) and the F 8801-B Meiosis project (Grant-DOI 10.55776/F88) of the Austrian Science Fund (FWF). FM is supported by ESP 566 (Grant-DOI 10.55776/ESP566) of the FWF. This research was funded in whole, or in part, by the FWF. Research at the IMP is supported by Boehringer Ingelheim. For the purpose of open access, the authors have applied a CC BY public copyright license to any Author Accepted Manuscript version arising from this submission.

## AUTHOR CONTRIBUTIONS STATEMENT

PB conducted all crosslink experiments, LC-MS measurements and wrote the manuscript. LT provided K562 cells and generated fluorescence images. FM contributed to 3D model prediction and interpretation. MM performed data analysis, post processing, interpretation and wrote the manuscript. JB, KM and MM conceptualized the study. All authors revised and agreed on the paper.

## DATA AVAILABILITY

The mass spectrometry raw proteomics data, Proteome Discoverer search results and used fasta files have been deposited to the ProteomeXchange Consortium via the PRIDE^28^ partner repository with the dataset identifier PXD061173. MS Amanda 3.0 (https://github.com/hgb-bin-proteomics/MSAmanda) and MS Annika 3.0, related workflow templates used for data analysis (https://github.com/hgb-bin-proteomics/MSAnnika) as well as code to generate XiView compatible exports from Annika data (https://github.com/hgb-bin-proteomics/MSAnnika_exporters/blob/develop/xiViewExporter_msannika.py are available on GitHub.

## REFERENCES

1. Iacobucci, C., Götze, M. & Sinz, A. Cross-linking/mass spectrometry to get a closer view on protein interaction networks. Curr. Opin. Biotechnol. 63, 48–53 (2020).

2. Botticelli, L. et al. Chemical cross-linking and mass spectrometry enabled systems-level structural biology. Curr. Opin. Struct. Biol. 87, 102872 (2024).

3. Sinz, A. Divide and conquer: cleavable cross-linkers to study protein conformation and protein–protein interactions. Anal. Bioanal. Chem. 409, 33–44 (2017).

4. Steigenberger, B., Albanese, P., Heck, A. J. R. & Scheltema, R. A. To Cleave or Not To Cleave in XL-MS? J. Am. Soc. Mass Spectrom. (2019) doi:10.1021/jasms.9b00085.

5. Müller, F. et al. A journey towards developing a new cleavable crosslinker reagent for in-cell crosslinking. 2024.11.05.621843 Preprint at 10.1101/2024.11.05.621843 (2024).

6. Burke, A. M. et al. Synthesis of two new enrichable and MS-cleavable cross-linkers to define protein–protein interactions by mass spectrometry. Org Biomol Chem 13, 5030–5037 (2015).

7. Kaake, R. M. et al. A New in Vivo Cross-linking Mass Spectrometry Platform to Define Protein–Protein Interactions in Living Cells. Mol. Cell. Proteomics 13, 3533–3543 (2014).

8. Matzinger, M., Kandioller, W., Doppler, P., Heiss, E. H. & Mechtler, K. Fast and Highly Efficient Affinity Enrichment of Azide-A-DSBSO Cross-Linked Peptides. J. Proteome Res. 19, 2071–2079 (2020).

9. Jiao, F. et al. DSBSO-Based XL-MS Analysis of Breast Cancer PDX Tissues to Delineate Protein Interaction Network in Clinical Samples. J. Proteome Res. 23, 3269–3279 (2024).

10. Zhang, H. et al. DDX39B contributes to the proliferation of colorectal cancer through direct binding to CDK6/CCND1. Cell Death Discov. 8, 1–9 (2022).

11. Zhao, G. et al. DDX39B drives colorectal cancer progression by promoting the stability and nuclear translocation of PKM2. Signal Transduct. Target. Ther. 7, 1–15 (2022).

12. Leitner, A. et al. Expanding the Chemical Cross-Linking Toolbox by the Use of Multiple Proteases and Enrichment by Size Exclusion Chromatography. Mol. Cell. Proteomics 11, M111.014126 (2012).

13. Hohmann, U. et al. A molecular switch orchestrates the nuclear export of human messenger RNA. 2024.03.24.586400 Preprint at 10.1101/2024.03.24.586400 (2024).

14. Abramson, J. et al. Accurate structure prediction of biomolecular interactions with AlphaFold 3. Nature 630, 493–500 (2024).

15. Fujita, K. et al. Structural differences between the closely related RNA helicases, UAP56 and URH49, fashion distinct functional apo-complexes. Nat. Commun. 15, 455 (2024).

16. Szklarczyk, D. et al. STRING v11: protein-protein association networks with increased coverage, supporting functional discovery in genome-wide experimental datasets. Nucleic Acids Res. 47, D607–D613 (2019).

17. Fasci, D., van Ingen, H., Scheltema, R. A. & Heck, A. J. R. Histone Interaction Landscapes Visualized by Crosslinking Mass Spectrometry in Intact Cell Nuclei. Mol. Cell. Proteomics 17, 2018–2033 (2018).

18. Fürsch, J., Kammer, K.-M., Kreft, S. G., Beck, M. & Stengel, F. Proteome-Wide Structural Probing of Low-Abundant Protein Interactions by Cross-Linking Mass Spectrometry. Anal. Chem. 92, 4016–4022 (2020).

19. Deng, W., Shi, X., Tjian, R., Lionnet, T. & Singer, R. H. CASFISH: CRISPR/Cas9-mediated in situ labeling of genomic loci in fixed cells. Proc. Natl. Acad. Sci. 112, 11870–11875 (2015).

20. Dorfer, V. et al. MS Amanda, a Universal Identification Algorithm Optimized for High Accuracy Tandem Mass Spectra. J. Proteome Res. 13, 3679–3684 (2014).

21. Birklbauer, M. J. et al. Proteome-wide non-cleavable crosslink identification with MS Annika 3.0 reveals the structure of the C. elegans Box C/D complex. Commun. Chem. 7, 1–17 (2024).

22. Doblmann, J. et al. apQuant: Accurate Label-Free Quantification by Quality Filtering. J. Proteome Res. 18, 535–541 (2019).

23. Hulsen, T. DeepVenn -- a web application for the creation of area-proportional Venn diagrams using the deep learning framework Tensorflow.js. Preprint at 10.48550/arXiv.2210.04597 (2022).

24. Combe, C. W., Graham, M., Kolbowski, L., Fischer, L. & Rappsilber, J. xiVIEW: Visualisation of Crosslinking Mass Spectrometry Data. J. Mol. Biol. 436, 168656 (2024).

25. Jumper, J. et al. Highly accurate protein structure prediction with AlphaFold. Nature 596, 583–589 (2021).

26. Mirdita, M. et al. ColabFold: making protein folding accessible to all. Nat. Methods 19, 679–682 (2022).

27. Yugandhar, K. et al. MaXLinker: Proteome-wide Cross-link Identifications with High Specificity and Sensitivity. Mol. Cell. Proteomics MCP 19, 554–568 (2020).

28. Perez-Riverol, Y. et al. The PRIDE database and related tools and resources in 2019: improving support for quantification data. Nucleic Acids Res. 47, D442–D450 (2019).

