## Supplemental Figures for "*In vivo* crosslinking and effective 2D enrichment for interactome studies of the nucleosome"

Phillip Bräuer[1], Laszlo Tirian[3], Fränze Müller[1], Julius Brennecke[3], Karl Mechtler[1,2,3, §], Manuel Matzinger[1,§]

[1] Research Institute of Molecular Pathology (IMP), Vienna BioCenter (VBC), Vienna, Austria.
[2] Gregor Mendel Institute of Molecular Plant Biology (GMI), Austrian Academy of Sciences, Vienna BioCenter (VBC), Vienna, Austria.
[3] Institute of Molecular Biotechnology (IMBA), Austrian Academy of Sciences, Vienna BioCenter (VBC), Vienna, Austria.

* These authors contributed equally

Supplemental Figure 1: Benchmarking of Sepharose vs Magnetic bead material

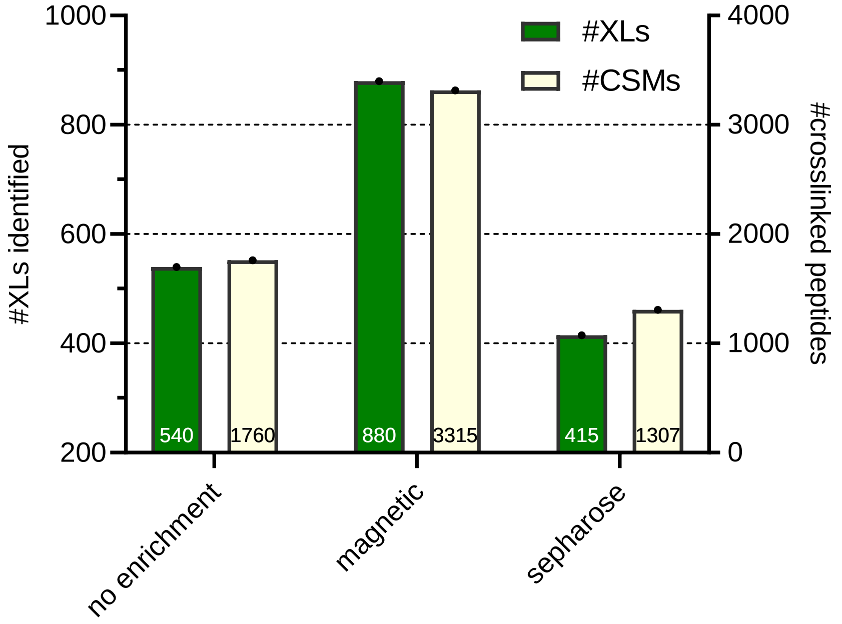

*20 µg DSBSO crosslinked Cas9-Halo peptides were either enriched using Sepharose or magnetic beads (both Cytivia) using 30µL bead slurry and the exact same processing workflow. 800 ng of each enriched eluate or of a non-enriched control sample were subjected to LC-MS analysis using a 2 h active gradient on an Orbitrap Exploris. Data was searched against a database containing 117 proteins (Cas9 + crapome). Bars indicate identified unique crosslink sites or crosslink sequence matches at 1% FDR level, n=1.*

Supplemental Figure 2: Distribution of crosslinked & monolinked peptides along the SEC gradient:

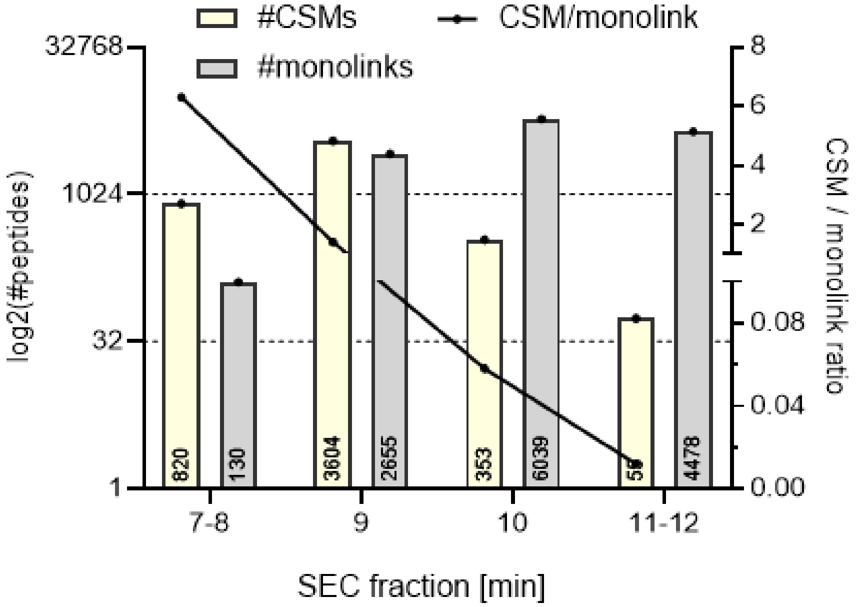

*Data from Figure 2 D-F, representative replicate, whole K562 cells were DSBSO linked and enriched using magnetic DBCO beads followed by SEC. Fractions from minute 7-12 were analyzed by means of LC-MS. Data was searched against the human proteome. Bars indicate identified crosslinked peptides (CSM) or monolinked peptides 1% FDR level, the line shows their ratio, n=1.*

Supplemental Figure 3: Abundance distribution of crosslinked proteins

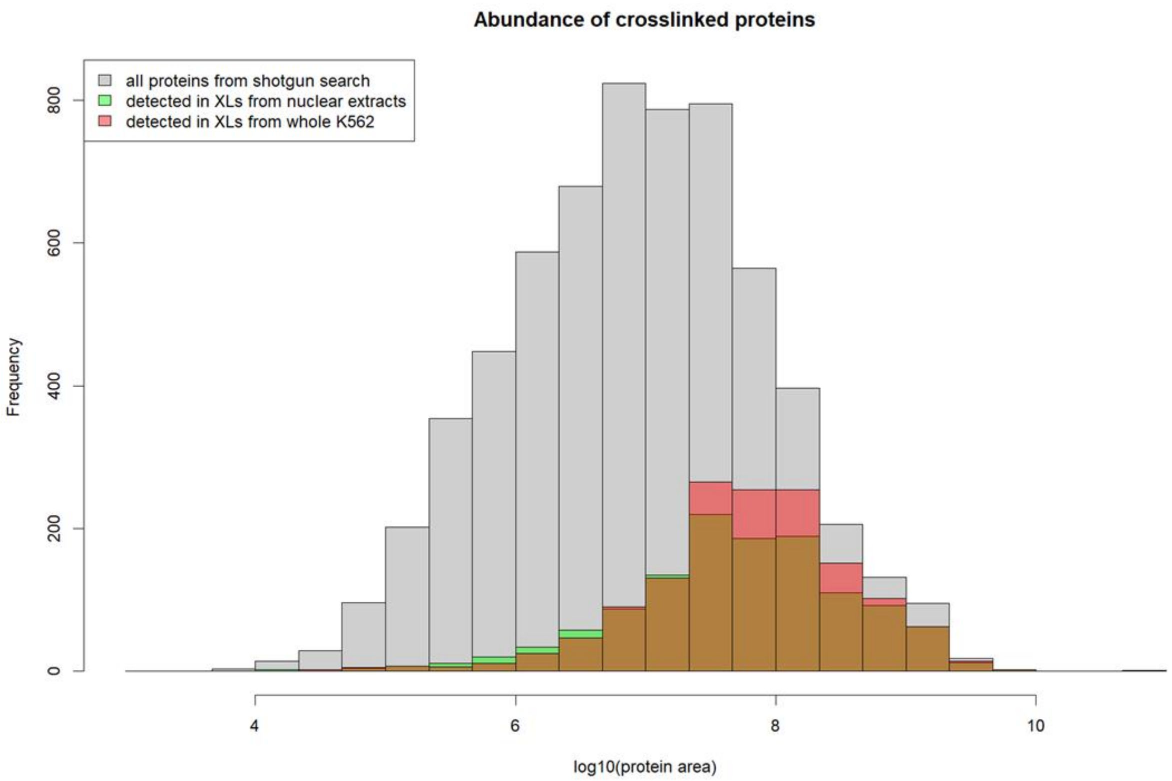

*Histogram of log transformed relative abundance of all proteins found within n=5 replicates of a shotgun analysis from K562 matched to those proteins crosslinked within n=3 replicates of our nuclear extract or whole cell samples when searching them towards the database created from that shotgun analysis as shown in Figure 3 C & D.*

Supplemental Figure 4: PPI network from whole K562 cells.

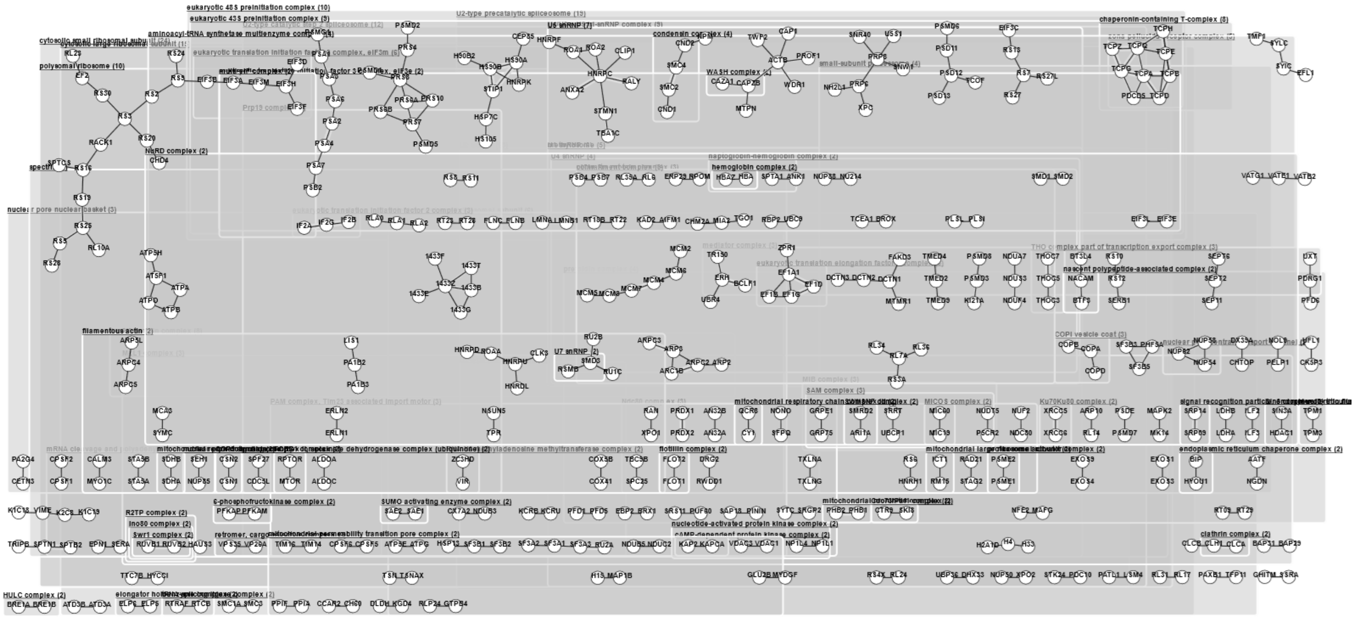

*PPI network as found from intact cell crosslinking samples after analysis against a shotgun database (see Figure 3C) with correlated groups annotated. In total 311 non-ambiguous PPI from 610 heteromeric crosslinks are shown.*
